# XAV-19, a swine glyco-humanized polyclonal antibody against SARS-CoV-2 Spike receptor-binding domain, targets multiple epitopes and broadly neutralizes variants

**DOI:** 10.1101/2021.04.02.437747

**Authors:** Bernard Vanhove, Stéphane Marot, Ray T. So, Benjamin Gaborit, Gwénaëlle Evanno, Isabelle Malet, Guillaume Lafrogne, Edwige Mevel, Carine Ciron, Pierre-Joseph Royer, Elsa Lheriteau, François Raffi, Roberto Bruzzone, Chris Ka Pun Mok, Odile Duvaux, Anne-Geneviève Marcelin, Vincent Calvez

**Affiliations:** Xenothera, Nantes, France; Sorbonne Université, INSERM 1136, Institut Pierre Louis d’Epidémiologie et de Santé Publique (iPLESP), Assistance Publique-Hôpitaux de Paris (AP-HP), Pitié Salpêtrière Hospital, Department of Virology, Paris, France; HKU-Pasteur Research Pole, School of Public Health, LKS Faculty of Medicine, The University of Hong Kong, Hong Kong SAR, P.R. China; Department of Infectious Disease, Nantes University Hospital, Nantes, France; INSERM CIC1413, Nantes University Hospital, Nantes, France; Department of Cell Biology and Infection, Institut Pasteur, Paris, France; Unité des Virus Émergents, UVE: Aix Marseille Univ, IRD 190, INSERM 1207, Hôpitaux Universitaires de Marseille, Marseille, France; Li Ka Shing Institute of Health Sciences, Faculty of Medicine, The Chinese University of Hong Kong, Shatin, Hong Kong; The Jockey Club School of Public Health and Primary Care, The Chinese University of Hong Kong, Hong Kong Special Administrative Region, China

**Keywords:** Polyclonal antibodies, pig, Covid-19, SARS-CoV-2, Spike, neutralization, variant of concern

## Abstract

Amino acid substitutions and deletions in Spike protein of the severe acute respiratory syndrome coronavirus 2 (SARS-CoV-2) variants can reduce the effectiveness of monoclonal antibodies (mAbs). In contrast, heterologous polyclonal antibodies raised against S protein, through the recognition of multiple target epitopes, have the potential to maintain neutralization capacities. XAV-19 is a swine glyco-humanized polyclonal neutralizing antibody raised against the receptor binding domain (RBD) of the Wuhan-Hu-1 Spike protein of SARS-CoV-2. XAV-19 target epitopes were found distributed all over the RBD and particularly cover the receptor binding motives (RBM), in direct contact sites with the Angiotensin Converting Enzyme-2 (ACE-2). Therefore, in Spike/ACE2 interaction assays, XAV-19 showed potent neutralization capacities of the original Wuhan Spike and of the United Kingdom (Alpha/B.1.1.7) and South African (Beta/B.1.351) variants. These results were confirmed by cytopathogenic assays using Vero E6 and live virus variants including the Brazil (Gamma/P.1) and the Indian (Delta/B.1.617.2) variants. In a selective pressure study with the Beta strain on Vero E6 cells conducted over 1 month, no mutation was associated with addition of increasing doses XAV-19. The potential to reduce viral load in lungs was confirmed in a human ACE2 transduced mouse model. XAV-19 is currently evaluated in patients hospitalized for COVID-19-induced moderate pneumonia in a phase 2a-2b (NCT04453384) where safety was already demonstrated and in an ongoing 2/3 trial (NCT04928430) to evaluate the efficacy and safety of XAV-19 in patients with moderate-to-severe COVID-19. Owing to its polyclonal nature and its glyco-humanization, XAV-19 may provide a novel safe and effective therapeutic tool to mitigate the severity of coronavirus disease 2019 (Covid-19) including the different variants of concern identified so far.

## Introduction

Passive antibody therapies have demonstrated efficacy to reduce progression of mild coronavirus disease 2019 (COVID-19) to severe disease if administered early enough in the course of illness ^1-3^. Three sources of antibodies have so far been assessed. First, passive antibody therapy using the infusion of convalescent plasma (CP) with high SARS-CoV-2 antibody titers in hospitalized patients, administered within 72 hours after the onset of mild symptoms, reduced the relative risk of progression to severe disease by 73% if CP presented a titer of >1:3200 and by 31.4% with lower titer CP ^1^. This was true with CP drawn between June and October 2020. However, the CP from patients infected by the original SARS-CoV-2 lineage had poor activity against the Beta variant and this was attributed to three mutations (K417N, E484K, N501Y) in the Spike protein ^4^. Among these mutations, E484K has been shown to play a major role to reduce the binding and neutralization ^5^. Second, beside use of CP, more than 50 neutralizing monoclonal antibodies (mAbs) are in development against the Spike receptor-binding domain (RBD) and the N-terminal domain (NTD) of SARS-CoV-2 ^6^. Those developed by Regeneron Pharmaceuticals (REGN-COV2 (casirivimab/imdevimab cocktail), Eli Lilly (bamlanivimab/etesevimab), Celltrion (regdanvimab) and GSK (sotrovimab) provide protection against the risk of severe COVID-19 when administered early in high-risk symptomatic patients with mild to moderate COVID-19 not requiring hospitalization ^7^. However, viral mutations can escape the mAbs which are used to treat the infection of SARS-CoV-2 ^8^. The Alpha variant is refractory to neutralization by most mAbs which target the NTD of Spike protein and also resistant to several RBD-specific mAbs ^9^. Mutations in the Beta lineage (K417N, E484K, and N501Y in RBD), especially mutations of Spike at E484 but also in the N-terminal Domain (NTD; L18F, D80A, D215G, Δ242-244, and R246I in SA variant ^10,11^) reduce neutralization sensitivity or confer neutralization escape from multiple mAbs ^4,5,12-20^. Third, polyclonal antibodies produced in their Fab’2 format from horses^21^ or in their IgG format from humanized cows ^22^ or glyco-humanized pigs ^23^ have also proven efficacy to neutralize SARS-CoV-2. The safety and tolerability in humans of Fab’2 from horses and of humanized IgG polyclonal antibodies has been confirmed recently in different clinical trials (Lopardo et al, 2021 ^21^, NCT04453384, NCT04469179, Gaborit et al, 2021 ^24^), contrasting with unmodified polyclonal antibodies containing wild-type IgG antibodies that induce serum sickness and allergic reactions (including fever and skin rashes) in 20 to 30% of the patients, excepting for those who concomitantly receive immunosuppression and high doses steroids ^25,26^. A partial efficacy of anti-SARS-CoV-2 Fab’2 from horses has been reported in those patients with negative baseline antibodies (NCT04494984). The efficacy of humanized or glyco-humanized IgG polyclonal antibodies in COVID-19 is still being investigated (NCT04453384, NCT04928430).

The possible advantage of polyclonal antibodies over mAbs is their recognition of an array of epitopes on the target antigen which should theoretically be not or less affected by antigen variations. Here, we investigated the extent to which XAV-19, a glyco-humanized swine polyclonal antibody previously shown to present neutralizing activity against SARS-CoV-2 Wuhan and D614G (B.1, PANGOLIN lineage) viruses ^23^, reduces viral load in vivo in human ACE2-expressing mice, binds multiple target epitopes on SARS-CoV-2 Spike and whether it maintains activity against the United Kingdom (Alpha/B.1.1.7), South African (Beta/B.1.351), Brazil (Gamma/P.1) and Indian (Delta/B.1.617.2) variants of concern.

## Methods

### Reagents

XAV-19 is a swine glyco-humanized polyclonal antibody against SARS-CoV-2 obtained by immunization of pigs with double knock out for alpha 1,3-galactosyltransferase (GGTA1) and cytidine monophosphate N-acetyl hydroxylase (CMAH) genes, as previously described ^23^. Intermediate R&D preparations of swine glyco-humanized polyclonal antibody against SARS-CoV-2 had been generated, presenting variable anti-SARS-CoV-2 binding activities ^23^. XAV-19 batches used in this study were clinical batches BMG170-B02, B03 and B06 which showed comparability in release testing. Comparator bamlanivimab is from Lilly (Indianapolis, In, USA). Recombinant Spike molecules of the Wuhan type (Sino Biological ref 40591-V08H), mutation-containing RBD (Y453F ref 40592-V08H80; N501Y, ref 40592-V08H82; N439K, ref 40592-V08H14; E484K, ref 40592-V08H84), Alpha (ref 40591-V08H12; containing mutations HV69-70 deletion, Y144 deletion, N501Y, A570D, D614G, P681H) and Beta (ref 40591-V08H10; containing mutations K417N, E484K, N501Y, D614G) forms and recombinant human Fc-tagged ACE-2 were purchased by Sino Biological Europe, Eschborn, Germany.

SARS-CoV-2 Wuhan (D614 and D614G B.1variant), Alpha, Beta, Gamma and Delta strains were isolated from SARS-CoV-2 infected patients in the Pitié-Salpêtrière, Aix-Marseille and Toulouse University hospitals (France). The BetaCoV/Hong Kong/VM20001061/2020 [HK1] Wuhan was isolated at The Chinese University of Hong Kong (China).

### Binding ELISA

The target antigen (SARS-CoV-2 Spike RBD-HIS protein, Sino Biological Europe) was immobilized on Maxisorp plates at 1µg/ml in carbonate/bicarbonate buffer at 4°C overnight. After washing, saturation was performed with PBS-Tween-BSA for 2h at room temperature. Samples were diluted into PBS-Tween and added into the plate in duplicate, incubated 2h at RT and washed 3 times. Bound pig IgGs were revealed with a secondary anti-pig-HRP-conjugated antibody (Bethyl Laboratories, USA) diluted in washing buffer, at 1:1000, incubated 1h at RT and washed 3 times. TMB reagent was added in the plate, incubated up to 20 minutes in the dark and the reaction was stopped with H_2_SO_4_. Reading was performed at 450nm.

### Spike/ACE-2 neutralization assay

An assay was developed to assess the properties of anti-SARS-CoV-2 Spike antibodies to inhibit binding of ACE-2 to immobilized Spike. SARS-CoV-2 Spike S1 (either Wuhan, Alpha or Beta) was immobilized on Maxisorp plates at 1 µg/ml in carbonate/bicarbonate buffer pH 9.0 at 4°C overnight. The plates were washed in PBS-Tween-0.05% and saturated with PBS-Tween-0.05%-2% skimmed milk for 2h at room temperature (RT). Anti-Spike RBD antibodies diluted in PBS-Tween-0.05%-1% skimmed milk were then added and incubated for 30 min. Then, ligand human ACE-2-mFc tag (Sino Biological; 125 ng/ml final concentration) was added in the same dilution buffer. After 1h incubation at room temperature and 3 washes, the mouse Fc tag was revealed with a specific HRP-conjugated anti-mouse IgG secondary antibody (diluted in in PBS-Tween-0.05%-1% skimmed milk powder at 1:1000, incubated 1h at RT and washed 3 times). TMB reagent was added into the plate, incubated 6 minutes in the dark and the reaction was stopped with 50 µl 1M H_2_SO_4_. The plate was read at 450 nm.

### Determination of XAV-19 target epitopes

A peptide microarray analysis has been performed using the PepStar™ system (JPT Peptide Technologies, Berlin, Germany). Fifty-three purified synthetic 15-meric overlapping peptides derived from the SARS-CoV-2 Spike S1 RBD domain (sequence from BDSOURCE accession number NC_045512.2), with an additional C-terminal glycine (added for technical reasons) were covalently immobilized on glass surface. Full-length human IgG, mouse IgG and pre-immune pig IgG were co-immobilized on microarray slides as assay controls. XAV-19 sample used in the analysis is the clinical drug substance batch BMG170-B06, hybridized at the dilution of 100 µg/ml (the same applied for control swine IgG) for 1 hour at 30°C on microarray slides. After sample incubation, secondary fluorescently labeled mouse anti-pig-IgG antibody diluted 1:5000 was added in the corresponding wells and left to react for 1 hour. Finally, a tertiary fluorescently labeled anti-mouse-IgG antibody at 1 μg/ml was incubated for 1 hour to detect bound anti-pig-IgG secondary antibody. After washing and drying, the slides were scanned with a high-resolution laser scanner at 635 nm to obtain fluorescence intensity profiles. The slides were scanned with a receiver gain of 900 V and images quantified to yield a mean pixel value for each peptide.

The epitope mapping process comprised a proteolytic digestion step of the SARS-CoV-2 Spike RBD protein, the isolation of resulting peptides by XAV-19 affinity chromatography and the LC-MS/MS analysis of the eluted peptides. In short, the RBD protein was reduced, alkylated and digested with an enzyme/protein ration of 1:50 during 3h at 37°C with the endopeptidases trypsin, chymotrypsin or Arg-C. Digestion products were immunocaptured on Sepharose 4B on which XAV-19 IgG has been immobilized, during 2h at room temperature. Binding yields on Sepharose was 80%. Columns were then washed (ammonium bicarbonate 25mM) and elution performed with a Glycine/HCl 50 mM pH2 buffer. Eluted fractions were then resolved by C18 inverted phase chromatography and tandem analyzed (MS/MS) to measure peptide masses. Data were compared with theoretical masses resulting from an in silico RBD digestion with the corresponding enzyme.

### Cytopathogenic Effect (CPE) assay

Vero cells (CCL-81) and Vero E6 cells (CRL-1586) were obtained from the American Type Culture Collection and maintained at 37°C with 5% CO_2_ in Dulbecco’s Modified Eagle’s Medium (DMEM), supplemented with 5% heat-inactivated fetal bovine serum (FBS) and 1X Penicillin-Streptomycin solution (Thermo Fisher Scientific, USA). SARS-CoV-2 clinical isolates (D614G variant; GenBank accession number MW322968), Alpha (GenBank accession number MW633280), Beta (GenBank accession number MW580244), Gamma (Gene accession number pending) and Delta (Gene accession number pending) were isolated from SARS-CoV-2 RT-PCR confirmed patients by inoculating Vero cells with sputum sample or nasopharyngeal swabs in the biosafety level-3 (BSL-3) facility of the Pitié-Salpêtrière University Hospital. Viral stocks were generated using one passage of isolates on Vero cells. Titration of viral stock was performed on Vero E6 by the limiting dilution assay allowing calculation of tissue culture infective dose 50% (TCID50). The neutralizing activity of XAV-19 was assessed with a whole virus replication assay using the five SARS-CoV-2 isolates. XAV-19 was subjected to serial two-fold dilution ranging from 50 µg/ml to 0.05 µg/ml in fresh medium. 50 µl of these dilutions were incubated with 50 µl of diluted virus (2 × 10^3^ TCID_50_/ml) per well in a 96-well plate at 37°C for 60 min in 8 replicates. Hundred µl of a Vero E6 cell suspension (3 × 10^5^ cells/ml) were then added to the mixture and incubated at 37°C under an atmosphere containing 5% CO_2_ until microscopy examination on day 4 to assess CPE. An infectivity score has been assigned on each well: 0, no cytopathic effect; 1, a fraction of cells was affected; 2, 100% cells affected. The addition of the scores in the 8 replicates was then transformed in percentage of the maximal scoring (ex. Score of 16 = 100%). For viral load (VL) quantification, a similar experiment was conducted with a range of XAV-19 dilution ranging from 24 µg/ml to 1 µg/ml in fresh medium. On day 4, RNA extraction of the 8 pooled replicates of each XAV-19 dilution was performed with NucliSENS EasyMag (BioMerieux) according to manufacturer’s protocol. The relative VLs were assessed from cycle threshold values for ORF1ab gene obtained by the TaqPath™COVID-19 RT-PCR (ThermoFisher, Waltham, USA) and by linear regression in log10 copies/ml with a standard curve realized from a SARS-CoV-2 positive nasopharyngeal sample quantified by Droplet-Digital PCR (Bio-Rad). IC50s were analyzed by nonlinear regression using a four-parameter dosage-response variable slope model with the GraphPad Prism 8.0.2 software (GraphPad Software, USA).To further analyze the neutralization potency and to confirm data with other independent laboratories, plaque reduction neutralization tests (PRNT)/ CPE assays and viral load evaluation were carried out independently on Vero E6 cells at the BSL-3 facility of VibioSphen, University Paul Sabatier, Toulouse, France and of Aix-Marseille University, Marseille, France. SARS-CoV-2 Wuhan, Alpha and Beta strains were isolated from patients with laboratory-confirmed COVID-19 from the corresponding University hospital. The viral isolates were amplified by one additional passage in VeroE6 cells to make working stocks of the virus. Vero E6 Cells were cultured in Dulbecco’s modified Eagle’s medium (DMEM) supplemented with 10% v/v fetal bovine serum, 1% v/v penicillin-streptomycin supplemented with 1% v/v sodium pyruvate at 1× 10^5^ cells per well in 12-well tissue culture plates. At 100% confluence (2 days post-seeding), the cells were washed twice with PBS and six serial dilutions of the virus (1/10 each time) were added to the cells. Following infection with 0.3 ml per well of each dilution, plates were incubated at 37°C for 1 h, and the cells were washed with PBS before the addition of 2% w/v agar containing 1 μg/ml-5 tosyl phenylalanyl chloromethyl ketone-trypsin (Sigma-Aldrich) to the cell surface. Plates were left at room temperature for 20–30 min to allow for the overlay to set and were then incubated at 37°C for 72 h. Cells were fixed with 4% v/v paraformaldehyde before both fixative and agar were removed, and cells stained with 0.1%w/v Crystal Violet (Fisher) in 20% v/v ethanol. Plaque titers were determined as plaque forming units per ml. CPE reduction assay was performed as follows: Vero E6 cells were seeded in 96-well clusters at a density of 5,000 cells/well 2 days before infection. Two-fold serial dilutions, starting from 100 µg/ml of XAV-19 were mixed with an equal volume of viral solution containing 300 pfu of SARS-CoV-2 (final volume 200 µL). The serum-virus mixture was incubated for 1 hour at 37°C in a humidified atmosphere with 5% CO_2_. After incubation, 100 µL of each dilution were added in 8 wells of a cell plate containing a semi-confluent Vero E6 cells monolayer. Control cells were infected with Covid-19 at MOI 0.01. Remdesivir (25 µM) was used as a positive control. After 3 days of incubation, the plates were inspected by an inverted optical microscope. Viable cells were quantified with CellTiter-Glo 2.0 luminescent cell viability assay.

### Antibodies escape study

XAV-19 was subjected to serial two-fold dilutions ranging from 50 µg/ml to 0.2 µg/ml in fresh medium. Fifty µl of XAV-19 dilutions were incubated with 50 µl of diluted virus (2 × 10^3^ TCID_50_/ml) per well in a 96-well plate at 37°C for 60 min in duplicates. Hundred µl of a Vero E6 cell suspension (3 × 10^5^ cells/ml) was then added to the mixture and incubated at 37°C under an atmosphere containing 5% CO2. A no antibody control was included to account for any cell culture adaptations of each SARS-CoV-2 variants. Virus replication was monitored on day 4 by screening for cytopathic effect. The supernatants were collected from wells with the highest antibody concentration displaying evident CPE. Fifty µl of these supernatants were passed under the same or greater fresh XAV-19 concentrations as before, until five passages. RNA extraction was also performed on these supernatants with NucliSENS EasyMag (BioMerieux) according to manufacturer’s protocol. Sanger sequencing was performed on the last passage (5^th^) of each variant.

### Human ACE-2 mouse model

The protocol of the animal experiments was described in the previous study ^27^. Balb/c mice were first infected with 10^8^ TCID_50_ of adenovirus carrying human ACE-2 protein intranasally. After 5 days post-infection, mice received intranasal administration of 10^5^ PFU of SARS-CoV-2 (BetaCoV/Hong Kong/VM20001061/2020 [HK1]). XAV-19 was administrated by intraperitoneal injection (I.P.) 24 hours before or after the infection. Lungs were collected at day 3 post-infection and viral load was measured by tissue culture infectious dose (TCID_50_) using Vero E6 cells. The animal experiments were performed in the BSL3 facility of the University of Hong Kong. The study protocol was carried out in strict accordance with the recommendations and was approved by the Committee on the Use of Live Animals in Teaching and Research of the University of Hong Kong (CULATR 5499-20).

## Results

### XAV-19 binding to SARS-CoV-2 Spike correlates with neutralizing potency

To assess whether binding strength to SARS-CoV-2 Spike of IgG antibodies present in XAV-19 is predictive of the neutralization potency, a series (n=117) of individual R&D serum samples drawn from immunized animals at different timepoints and presenting with different binding activities were evaluated in parallel in a Spike/ACE-2 interaction ELISA. The data (Figure 1A) indicated a global correlation (R=0.8) with high binders presenting with high ability to inhibit Spike/ACE-2 interaction. Then 4 R&D anti-SARS-CoV-2 IgG batches made from serum samples presenting variable anti-RBD binding activity levels were assessed in parallel in neutralizing ELISA and in CPE assays using live Wuhan viruses. The batches presented IC50 values by ELISA of 1.3, 1.34, 2.2 and 12 µg/ml and the corresponding values in TCID assays were of 3, 2, 12.5 and 25 µg/ml (R=0.91; Figure 1B).

**Figure 1.**
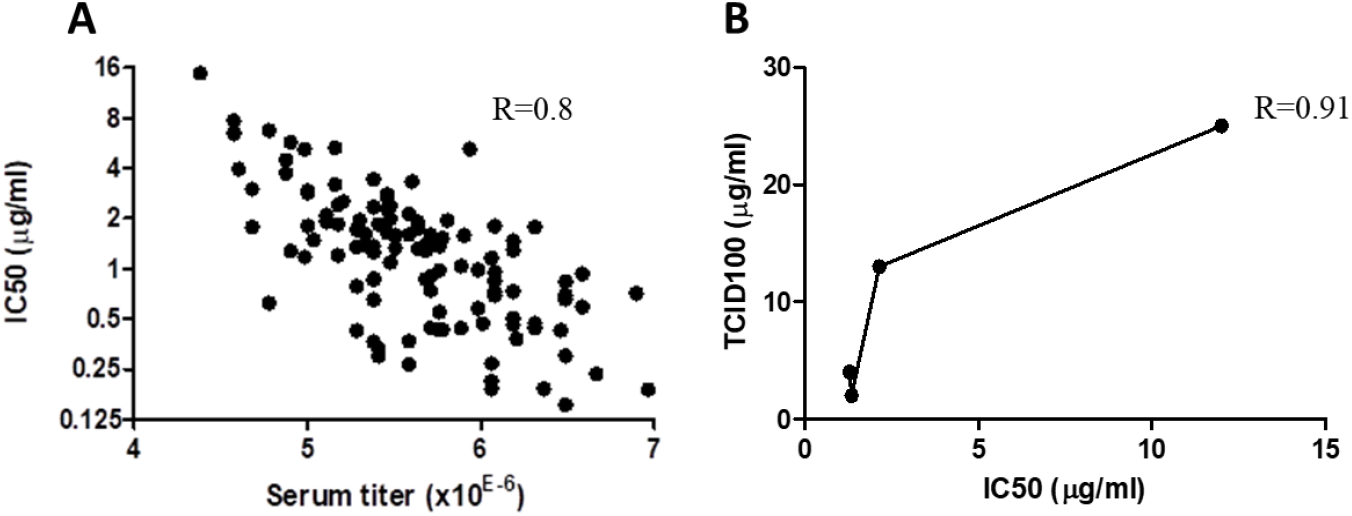
Correlation between binding, neutralizing ELISA, and CPE assay. A: a series of 117 individual hyperimmune serum samples were assessed in parallel in a SARS-CoV-2 Spike RBD binding ELISA and in a Spike/ACE-2 interaction assay. Linear correlation was observed with R= 0.8. Dots represent data from duplicate measurements in a single experiment. B: Four R&D batches of swine anti-SARS-CoV-2 Spike RBD polyclonal IgG were produced, with different binding activities against SARS-CoV-2 Spike ^23^. These samples were evaluated in parallel in the Spike/ACE-2 interaction assay, as in A, and by CPE. IC50 in ELISA and CPE, expressed in TCID_100_ are from single experiments and have been plotted to evaluate correlation. R value after linear extrapolation was 0.91.

### XAV-19 target epitopes on SARS-CoV-2 Spike protein

Two orthogonal methods have been used to identify the epitopes recognized by XAV-19 on the S1-RBD protein. First, all 15-meric peptides overlapping by 10 amino acids (thus a total of 53 peptides) were spotted on glass slides and hybridized with XAV-19 or pre-immune swine GH-pAb control antibodies. The resulting heatmap plot revealed that all peptides, though to varying extents, could be specifically recognized by antibodies contained in XAV-19 (Figure 2A). Since not all peptides can be linked together with XAV-19 antibodies when contained in the Spike protein, for steric hindrance, a second investigation was undertaken to identify which peptides in the Spike RBD domain in a more native configuration are recognized by XAV-19 antibodies. The assay was based on the recognition by XAV-19 antibodies of SARS-CoV-2 Spike RBD-derived peptides obtained by proteolytic digestion (three enzymes tested: trypsin, chymotrypsin and Arginase-C) and an isolating step of the resulting peptides by affinity chromatography (XAV-19 being immobilized on Sepharose) followed by an LC-MS/MS analysis of the eluted peptides. This epitope mapping analysis revealed several recognition areas on the S1-RBD protein. The peptides where the two methods gave the strongest overlapping hits, thus most probably representing dominant target epitopes, were amino acids 347-355 and 445-461. Amino acids 409-417, 462-473 and 530-535 were also found to be protected in the LC-MS/MS analysis although less recognized in the peptide array (Figure 2B). Interestingly, 6 amino acids described to directly interact with human ACE-2 ^28^ are located in these regions.

**Figure 2.**
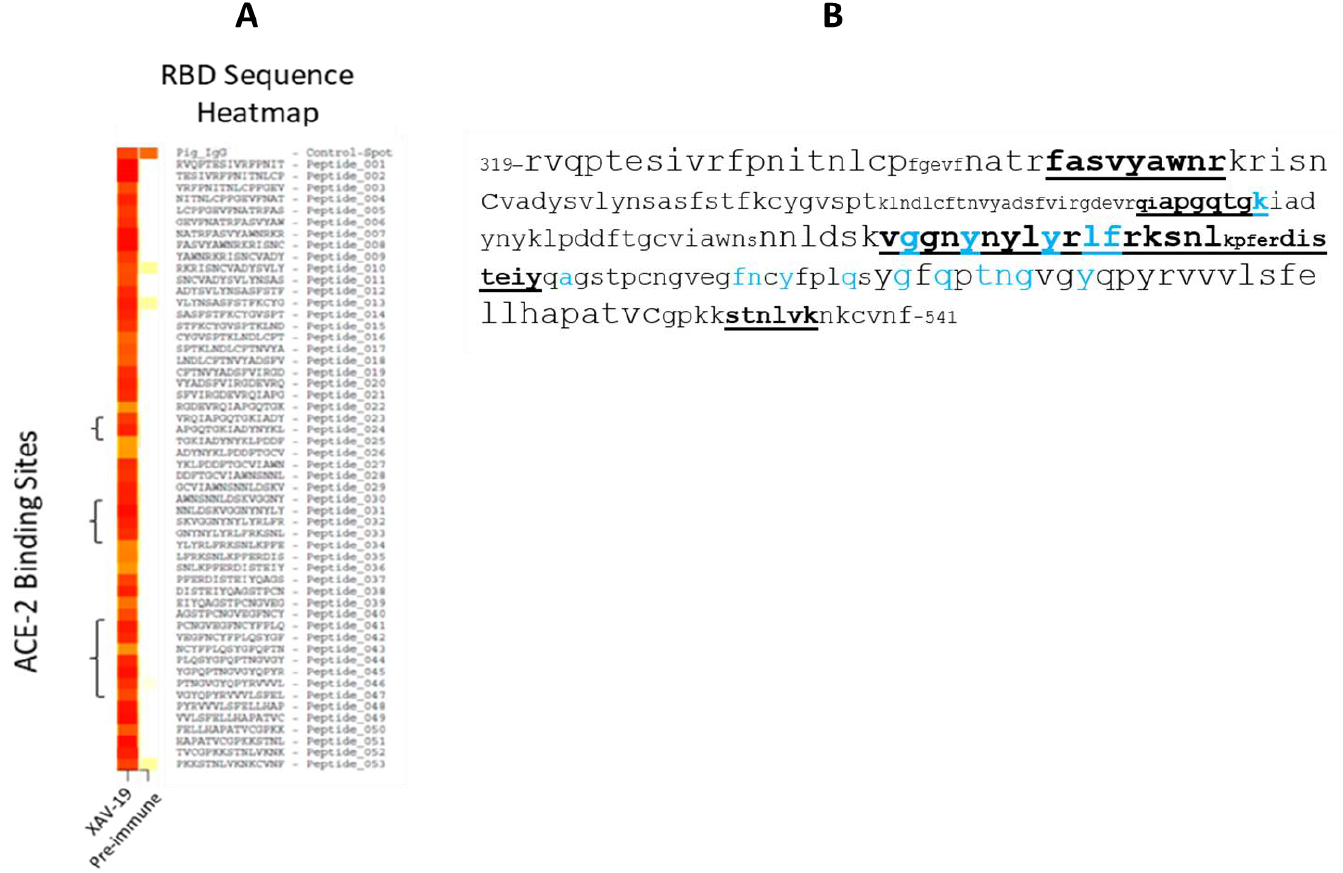
XAV-19 Target epitopes. A: heatmap from a peptide array experiment showing binding intensity of XAV-19 and pre-immune IgG, to 15-meric peptides of the SARS-CoV-2 Spike RBD sequence, overlapping by 10 amino acids. White, no binding; yellow, background binding; light red, medium binding; dark red, strong binding. B: XAV-19 target epitopes on SARS-CoV-2 Spike RBD domain (amino acid sequence numbered according to DBSOURCE sequence reference NC_045512.2) as recorded in LS-MS/MS peptide mapping. Font size refers to binding intensity in the peptide array shown in A: font 10, weak recognition; font 14, medium recognition; font 18, strong recognition. Underlined bold: XAV-19 target peptides as defined in LS-MS/MS. Blue characters: amino acids in contact with ACE-2, according to Jafary et al. ^28^.

### ELISA to assess inhibition of Spike from Wuhan, Alpha and Beta variants interaction with ACE-2

XAV-19 was tested in a Spike/ACE-2 binding competition assay, where the Spike protein was of the original Wuhan type or contained the RBD mutations N501Y, N439K, Y453F described in the Alpha and Beta variants, or the mutation E484K show to induce resistance to mAbs ^29^. Variants expressing a combination of mutations present in the Spike Alpha (HV69-70 deletion, Y144 deletion, N501Y, A570D, D614G, P681H) or Beta (K417N, E484K, N501Y, D614G) were also tested. All single mutation forms of the Spike could be fully neutralized at concentrations not significantly different (slightly lower for the E484K mutation) from the Wuhan type (Figure 3A). XAV-19 also demonstrated a 100% inhibitory activity on the 2 Spike proteins fully representative of the Alpha and Beta variants, similar to the Wuhan Spike, with IC50 values of 6.4, 4.0 and 4.5 µg/ml, respectively (Figure 3B). Bamlanivimab, tested in parallel, demonstrated a potent inhibitory capacity against the Wuhan and the Alpha variants, with a IC50 value of 0.01µg/ml but, as described ^30^, failed to inhibit binding of Beta Spike to ACE-2, even at high concentration (Figure 3C).

**Figure 3.**
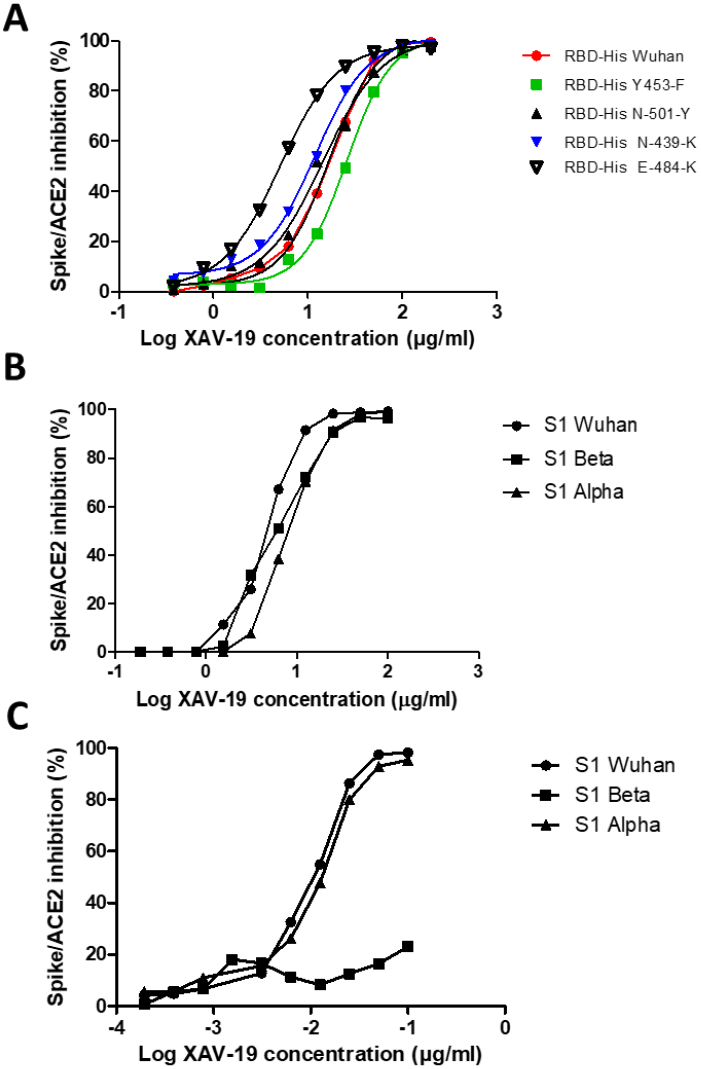
Neutralization assay in the ELISA format: assessment of SARS-CoV-2 Spike/ACE-2 interaction and its anti-RBD antibody-mediated inhibition. Spike-HIS containing the indicated mutations (A) or grouped mutations corresponding to the Alpha and Beta variants (B, C) was immobilized on plastic and binding of recombinant human ACE2-Fc was revealed with a secondary antibody against Fc. 100% inhibition represents absence of Spike/ACE-2 interaction. (A, B) Means of triplicate measurements run in a single experiment assessing XAV-19 at the indicated concentration. (C) Means of triplicate measurements run in a single experiment assessing bamlanivimab at the indicated concentration.

### Neutralization of live SARS-CoV-2 variants in CPE

The neutralizing effect of XAV-19 was determined by CPE assays in 3 different platforms (Paris Sorbonne university, Aix-Marseille university, Paul Sabatier Toulouse university) using Wuhan (D614), B.1 (D614G PANGOLIN lineage) (D614G), Alpha, Beta, Gamma and Delta SARS-CoV-2 clinical isolates, as previously described ^31^. The test assessed inhibition of live viruses with sensitive Vero E6 cells and recorded infection after 4 days by assessing CPE and viral load by RT-qPCR. Data showed similar neutralizing potency for the Wuhan B.1 (D614G), Alpha and Beta strains in a first set of experiments (Supplementary Figure 1) and showed global similar potency of XAV-19 on the Wuhan D614G, Alpha, Beta, Gamma and Delta strains in a second set of experiments (Figure 4), with absence of neutralizing activity below 1.5 µg/ml and 100% neutralizing activity above 5 to 10 µg/ml.

**Figure 4.**
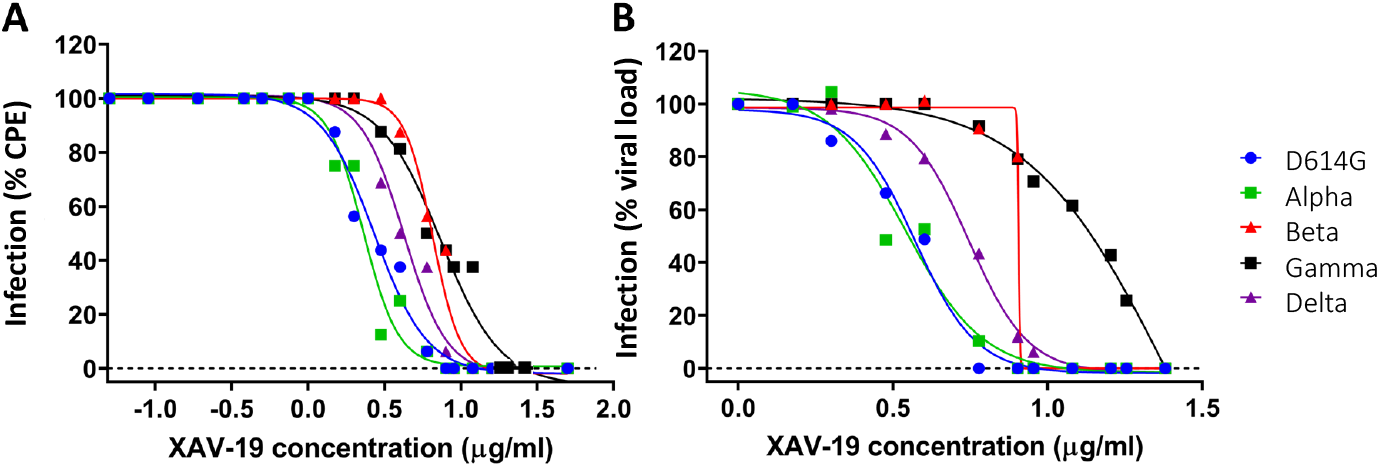
In vitro neutralization of SARS-CoV-2 variants by XAV-19. XAV-19 neutralizing potency was evaluated in an in vitro assay using whole replicating viruses. Percentage of infection was evaluated as described in Materials and Methods, based on cytopathogenic effect (CPE) (A) and virus RNA load (B) after infection with SARS-CoV-2 viruses of the indicated variants. CPE percentage was assessed by microscopy examination and calculated on 8 replicates for each XAV-19 concentration. 100% represent absence of CPE inhibition at the studied concentration, as found in control (no inhibitor) condition. Viral load percentage was calculated as the ratio of viral load in each XAV-19 concentration to viral load in controls (no inhibitor). XAV-19 concentrations are expressed on a 10 logarithmic scale. Blue dot: D614G/B.1 variant; Green square: Alpha/UK/B.1.1.7 variant; Red triangle: Beta/SA/B.1.351 variant; Black square: Gamma/BR/P.1 variant; Purple triangle: Delta/B.1.617.2 variant.

IC50 to reduce CPE were as follows, different values for a single strain reflecting experiments performed independently in different labs: for Wuhan, 0.4 µg/ml, B.1 D614G, 2.2/2.7 µg/ml; Alpha, 0.1/2.2/2.3 µg/ml; Beta, 3.2/6.5 µg/ml; Gamma, 8.1 µg/ml; Delta 4.2 µg/ml. IC50 to reduce the viral load assessed by RT-qPCR were for Wuhan D614, 0.5/3.6 µg/ml; B.1 D614G, 0.5/3.6/3.7 µg/ml; Alpha, 1.2/3.6/3.4 µg/ml; Beta, 6/8.9/8.0 µg/ml; Gamma, 2.4/13.8 µg/ml; Delta 5.5 µg/ml.

### Absence of XAV-19 induced SARS-CoV-2 variant selection in vitro

Antibodies, by applying a selective selection pressure, can favor outgrowth of resistant novel variants with unknown properties, especially in conditions where their neutralizing potency is suboptimal. To investigate whether XAV-19 is susceptible to generate such variants, Vero E6 cells were infected with 100 TCID50/50 µl of either B.1D614G, Alpha, Beta, Gamma or Delta strains and maintained over 5 passages (20 days) with culture medium or culture medium containing increasing concentrations of XAV-19. After passage 5, emergence of antibodies escape mutants was evaluated by Sanger sequencing. The data (Table 1) show absence of mutations in the RBD domain (amino-acids 331 to 524) under any condition. Variations were found in the Spike outside the RBD domain, not associated with addition of XAV-19. Overall, 11 mutations were found without antibody addition (culture medium only), whereas 5 mutations were found with XAV-19. These mutations probably represent adaptations to the culture conditions. None of these mutations has been described as resistant to antibodies.

**Table 1:** Amino-acid substitutions in the SARS-CoV-2 Spike detected after 5 passages in Vero E6 cells with or without addition of XAV-19.

|  | **Bav Pat D614G** | **Alpha** | **Beta** | **Gamma** | **Delta** |
| --- | --- | --- | --- | --- | --- |
| **No XAV-19 control** | G72R/G,<br>D215N/DS1249F/S | S151R | H66R | ∅ | M153R/M<br>S686R |
| **XAV-19 increasing**<br>**concentration** | S1170F | S151R | H66H/R | S813I/S | ∅ |

### Reduction of viral load by XAV-19 in human ACE-2 expressing mice

Balb/c mice with human ACE-2 expression in the lung were infected with 10^5^ PFU SARS-CoV-2 intranasally. XAV-19 was administrated through intraperitoneal injection under different experimental conditions: 1) 20 mg/Kg, 24h before viral infection. 2) 2 mg/ kg, 24h before viral infection. 3) 0.2 mg/ kg, 24h before viral infection. 4) 20 mg/ kg, 24h after viral infection. 5) untreated. The results showed a 98% reduction of viral load in the lung at day 3 if 20 mg/kg of XAV-19 was given to the mice 24 hours before the infection and a 94% reduction if 20 mg/Kg of XAV-19 was given 24h after the infection. No significant reduction of viral load was observed from the group of administrating either 2 or 0.2 mg/kg 24 hours before infection (Figure 5A). No virus was found in all groups at day 5 of post-infection (Data not shown).

**Figure 5.**
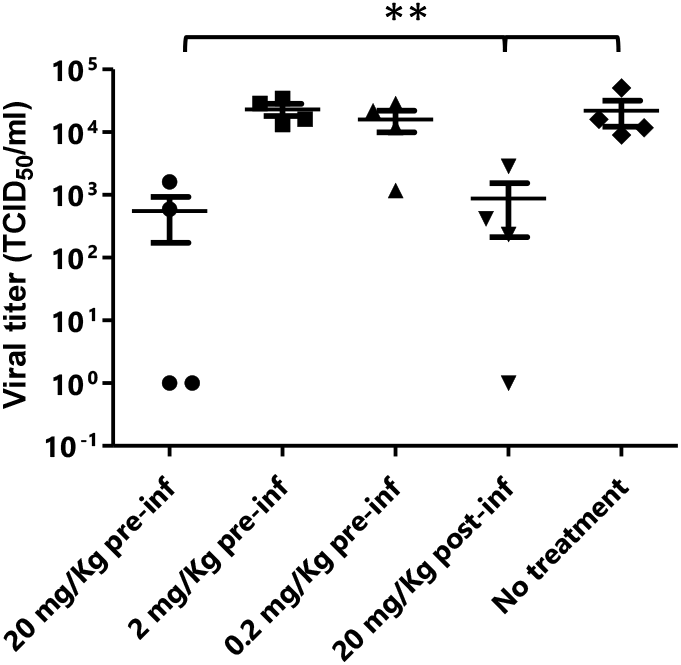
Murine model of SARS-CoV-2 infection. Balb/c mice (n=4/group) were first transduced intranasally with adenovirus coding for human ACE-2. After 5 days, mice were infected intranasally with SARS-CoV-2 (Wuhan strain) and received the indicated dose of XAV-19, intraperitoneally, 24h before (pre-inf) or after (post-inf) infection. Three days later, lungs were necropsied, homogenized and viral load was evaluated by PRNT. Each dot represents a single mouse and is the mean of 8 replicates assessed in a single experiment. Nonparametric statistics were used for pairwise comparisons using Kruskal-Wallis tests. **, p<0.01.

## Discussion

XAV-19 is a swine glyco-humanized neutralizing polyclonal antibody raised against the Spike RBD protein of the SARS-CoV-2 original Wuhan strain ^17^. In this paper, we report that XAV-19 also fully neutralizes Alpha (United Kingdom/B.1.1.7), Beta (South African/B.1.351), Gamma (Brazil/P.1) and Delta (Indian/B.1.617.2) variants in vitro. Because XAV-19 is manufactured after pooling many individual serum samples drawn on multiple donor animals, it presumably contains a diversity of IgG molecules presenting an array of avidities and seeing several epitopes on the target protein. When analyzing individual sera (from individual donor animals) composing XAV-19, a correlation was observed between binding and neutralizing activities. This indicated that globally, most antibodies in XAV-19 bind to SARS-CoV-2 RBD in a way that prevents SARS-CoV-2 Spike interaction with human ACE-2. Because the RBD antigen and the Spike-HIS protein used in neutralizing assays are recombinant molecules, it was important to control that the corresponding RBD domain in live viruses could also be similarly targeted and neutralized. Parallel testing of 4 XAV-19 R&D batches presenting with varying neutralization potencies confirmed that a high neutralization potential assessed in vitro with a recombinant Spike-Fc molecule corresponded to a high neutralization potential in cytopathic assays, using live viruses. This observation confirmed data showing that a neutralizing ELISA is predictive of SARS-Cov-2 neutralization assessed in a lentivirus-pseudotyped SARS-CoV-2 neutralization assay ^4^.

Target epitopes of XAV-19 on Spike RBD differed whether assessed by peptide microarray or by peptide mapping. Peptide microarrays indicated that all possible 15-meric peptides contained in the Spike RBD protein have been immunogenic in animals and generated specific antibodies. In real situation, however, not all antibodies can be engaged together and some of them might fail to get access to the corresponding protein domain, for steric hindrance. The peptide mapping assay allows presentation of large peptides issued from RBD proteolysis and therefore is believed to be more relevant. The peptide mapping partially confirmed data from the peptide microarrays and identified several target epitopes containing amino acids important for the interaction between Spike and human ACE-2, explaining the neutralizing activity of XAV-19. The Alpha, Beta, Gamma and Delta variants contain mutations in the Spike RBD domain, which might impact recognition by anti-RBD antibodies. K417N/T and E484K are present in Beta and Gamma variants, N501Y is present in the Alpha, Beta and Gamma variants and T478K appears in the Delta variant. These mutations are located in peptides scored as “medium” in peptide microarray, and not revealed by peptide mapping. These mutations might therefore have no or a limited impact on XAV-19 binding. In contrast, L452R is present in the Delta variant and located in a region seen as a major target epitope by both techniques. This predicts that antibodies binding to the corresponding peptide would be affected. However, XAV-19 behave similarly towards the Delta variant, with even a stronger neutralization of the viral load after infection of Vero E6 cells (Figure 4B). It is possible that in a polyclonal antibody, although a specific antibody is prevented from recognizing its target (because of a change in the epitope), other antibodies remain able to bind a closely related epitope. This is what is suggested by the peptide array data showing many binding possibilities.

Independent cytopathic assays with live viruses were run in parallel in different locations by different teams. They show similar findings, i.e., that a concentration of XAV-19 up to 10 µg/ml is required to fully neutralize all the variants. There was a clear tendency of higher concentration required to exhibit similar neutralizing capacity against the Beta and Gamma variants, as compared with Wuhan, Alpha and Delta forms. This difference was observed in different assays, whether measuring target cell viability, cytopathogenic effect or viral RNA load, and confirmed in different experiments, thus reflecting actual differences rather than experimental variability. Thus, our results demonstrate that XAV-19 can fully neutralize all SARS-CoV-2 variants, and a clear difference was evident when compared to bamlanivimab, which has no neutralizing effect on the Beta variant.

When administered intraperitoneally to human-ACE-2 mice challenged intranasally with SARS-CoV-2 viruses, XAV-19 induced a dose-dependent reduction of viral load in the lung, demonstrating the potential to localize to infected tissues. The 94% to 98% reduction in the viral load noticed here is in agreement with data obtained with different neutralizing antibodies in ferrets, hamster or mouse models ^32,33,34-40^ or obtained with the REGN-CoV2 antibody cocktail in a comparable animal model ^41^. In a few studies, however, the viral reduction factor reached 3 or 4 log ^33,42-45^. Interestingly, Gilliland et al ^22^ reported absence of viral reduction in the lung of SARS-CoV-2 challenged mice treated with a humanized cow polyclonal antibody, whereas a clear clinical impact was demonstrated. One limitation of our model in which mice airways are transduced with an adenovirus expressing human ACE-2 is the absence of clinical symptoms, as the mice eliminate the virus by themselves within a few days. The evaluation we made was restricted to the assessment of the viral load in lung. Therefore, owing to the lack of correlation between lung viral load and clinical status, our data cannot inform whether XAV-19 can bring a clinical benefit.

Early after Covid-19 outbreak onset, many labs have been able to rapidly develop neutralizing antibodies. One year later, as of mid-2021, more than 93 clinical trials assessing the safety and benefit of mAbs and 8 of polyclonal antibodies are listed in the Clinicaltrials.gov repository (NCT04610502, NCT04838821, NCT04514302, NCT04834908, NCT04834089, NCT04518410, NCT04453384, NCT04453384). In late 2020, variants of concern started to spread in the population and now cause the majority of infections. However, antibodies that are now assessed clinically have been mostly raised against the initial, Wuhan strain. It has therefore become essential to revisit their potential to also neutralize variants. XAV-19 is currently being tested in phase 2 and 3 studies (NCT04453384; Eudract Number: 2020-005979-12). The Phase 2a demonstrated that a single intravenous perfusion of XAV-19 at 2 mg/kg was safe, achieving a median serum C_max_ of 50.4 µg/ml and day 8 concentration of 20.3 µg/ml with a elimination half-life (T1/2) estimated at 11.4 days ^24,46^. The data presented here, together with these pharmacokinetic data, indicate that XAV-19 can provide high and sustained therapeutic activity in vivo. These data warrant continuation of clinical studies with XAV-19, especially in a context where the Delta variant becomes dominant and other variant of concerns emerge.

## Authorship

Conceived the study: OD, BV

Designed and supervised some experiments: OD, BV, RB, FR, FT, XDL, SM, AGM, VC

Performed the experiments: GE, RTS, AGM, CC, PJR, SM, IM, CKPM

Analyzed data: BV, OD, AGM, GL, EL, EM, VP, BG, FR

## Acknowledgments

We sincerely thank Prof. Xavier de Lamballerie and Dr. Franck Touret from UMR IRD 190, Inserm 1207 “Unité des Virus Émergents”, Aix-Marseille Université for their active technical contribution to generating neutralization data and for their advice. We also thank the virology departments of Saint-Antoine and Avicenne Universitary hospitals who gently shared us the clinical specimen allowing us to isolate the Gamma/P.1 and Delta/B.1.617.2 variants respectively. We also thank Prof Jincun Zhao and Yanqun Wang to provide us the adenovirus which carries the human ACE-2.

This work was supported by Xenothera, the Agence Nationale de la Recherche sur le SIDA et les Maladies Infectieuses Emergentes (ANRS MIE), AC43 Medical Virology and Emergen Program, the SARS-CoV-2 Program of the Faculty of Medicine of Sorbonne Université and by Bpifrance, grant « Projet de Recherche et Développement Structurant Pour la Compétitivité spécifique à la crise sanitaire COVD-19 – POLYCOR» and the National Research Foundation of Korea (NRF) grant funded through the Korea government (NRF-2018M3A9H4055203).

## Competing Interests

The authors of this manuscript have conflicts of interest to disclose: OD, PJR, CC, GE, EL, BV are employees of Xenothera, a company developing glycol-humanized polyclonal antibodies as those described in this manuscript.

## Notes

### Summary of Updates

Novel findings on target epitopes of XAV-19 on SARS-CoV-2 spike RBD have been added, as well as neutralizing capacities against the delta variant and in vivo mice experiments.

