## Supplementary figures and images for "XAV-19, a swine glyco-humanized polyclonal antibody against SARS-CoV-2 Spike receptor-binding domain, targets multiple epitopes and broadly neutralizes variants"

### Supplementary Figure 1

Supplementary Figure 1

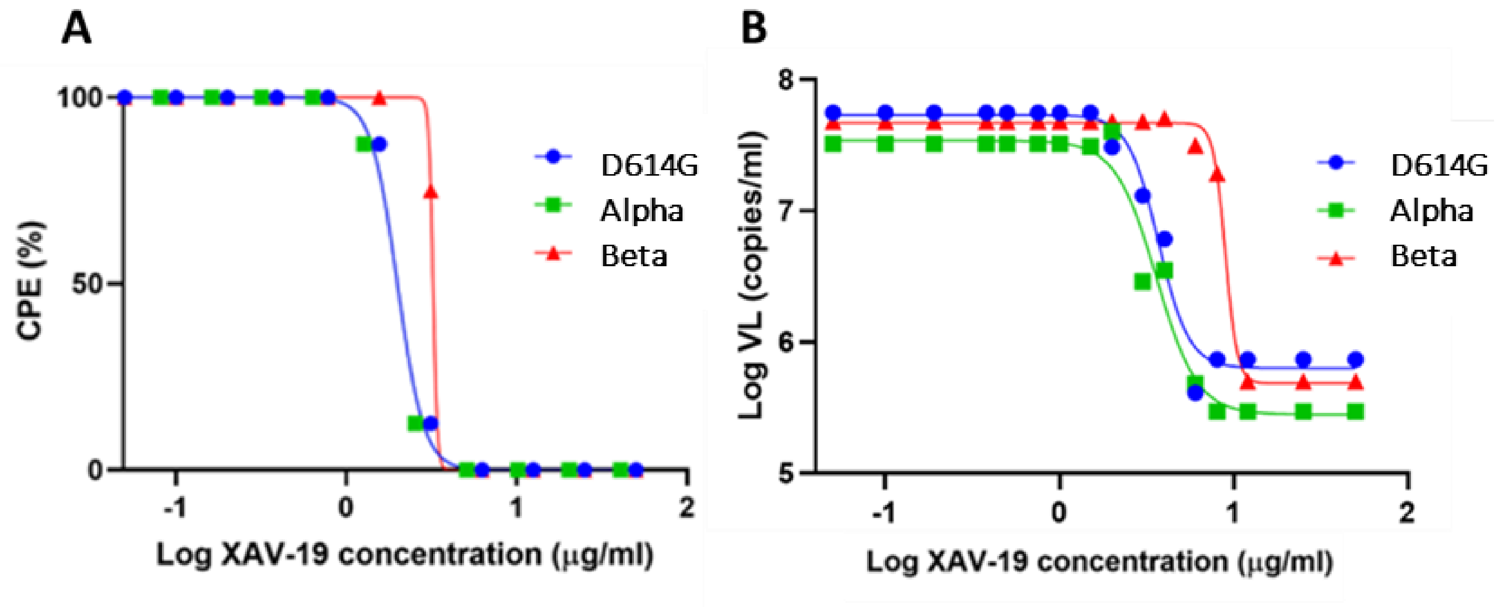
